# Pan-genotypic probe-based enrichment to improve efficiency of Hepatitis B virus sequencing

**DOI:** 10.1101/2023.02.20.529276

**Authors:** Sheila F Lumley, Daisy Jennings, Elizabeth Waddilove, Amy Trebes, Marion Delphin, Louise O Downs, George MacIntyre-Cockett, Yanxia Wu, Sandra Chaudron, Catherine de Lara, Haiting Chai, Tongai G Maponga, Jacqueline Martin, Jane Collier, Camilla LC Ip, Eleanor Barnes, David Bonsall, Paolo Piazza, M. Azim Ansari, Philippa C Matthews

**Author notes:** these authors contributed equally (Sheila F Lumley, Daisy Jennings, Elizabeth Waddilove). these authors contributed equally (M. Azim Ansari, Philippa C Matthews).

## Abstract

Hepatitis B Virus (HBV) genome sequencing can be used to provide more complete genetic information at the population and individual level to shed light on the limitations of current interventions, and inform new strategies for elimination. HBV sequencing is challenging due to the partially dsDNA genome, high diversity, low viral loads and presence of large amounts of host genetic material in clinical samples. Here we describe the design and use of a pan-genotypic panel of 74 HBV specific capture-probes and nuclease treatment in improving sequencing efficiency. We processed 20 plasma samples (viral loads 1.98 to 4.07 log_10_, genotypes A-E) and three positive controls (human total brain RNA and bacteriophage lambda DNA) in triplicate to compare DNAse vs. RNAse vs. no nuclease treatment. We prepared libraries using the Takara Bio SMARTer Stranded Total RNA-Seq Kit v3, split the library in two, enriching half with the custom-designed probe panel and xGen Hybridization and Wash Kit (IDT), the other half was not enriched. Both libraries were sequenced on the NovaSeq6000 platform with 2×150nt paired-end reads. Capture resulted in a 47,970 fold increase in the number of reads mapped to the HBV genome in the “no nuclease” arm (243 HBV reads per million reads sequenced in the capture pool vs. 5×10^−3^ reads per million in the no-capture pool). Out of 20 samples, only 1 without capture generated HBV reads (viral load 3.89 log_10_ IU/ml) vs. 19 samples with capture. HBV sequence yield was increased in the capture arm and resulted in 2.30 log_10_ (95% confidence interval 1.99 - 2.48 log_10_) increase in HBV reads (per million reads sequenced) per log_10_ increase in viral load. The proportion of HBV reads increased a median of 12 fold with RNAse treatment. We developed a targeted pan-genotypic sequencing method using a custom panel of biotinylated oligos that increases the sequencing efficacy of HBV. This method will allow us to gain a better insight into HBV diversity.

## INTRODUCTION

Chronic hepatitis B virus (HBV) infection is a significant public health burden with over 300 million infected individuals worldwide. Despite the availability of a prophylactic vaccine and suppressive antiviral therapy there are around 555,000 deaths each year due to HBV infection ^1^. HBV genome sequencing can be used to provide more complete genetic information at the population and individual level to shed light on the limitations of current interventions, and inform new strategies for elimination ^2,3^.

HBV has a circular, partially double-stranded (ds) DNA genome of approximately 3.2 kB (relaxed-circular (rc-DNA)). The virus replicates via a full length pre-genomic (pg)RNA intermediate, leading to a higher genome variability compared to other DNA viruses. Nine different HBV genotypes and one putative genotype (A–J) have been defined by >8% divergence at the nucleotide level ^3^. Sequencing is challenging due to the unusual genome structure, high diversity and low viral loads particularly in HBeAg negative individuals (median viral load of 3.2 log_10_ IU/ml) ^4^. The challenge of sequencing is exacerbated by the high proportion of host genetic material present in plasma samples. Strategies for increasing HBV yield broadly involve host depletion (e.g. nuclease treatment, filtration, ultracentrifugation ^5^), viral amplification (e.g. polymerase chain reaction ^6,7^, rolling circle amplification ^8,9^) or viral enrichment (CRISPR cas9-mediated enrichment, probe-based enrichment) ^10–14^.

Probe based enrichment has been used to improve the viral sequencing efficiency for other viral pathogens ^12,15–17^. Biotinylated single stranded DNA probes designed to hybridize to target viral sequences within a sequencing library selectively capture the target viral sequences. Here we describe the design and use of a pan-genotypic panel of probes to improve sequencing yield for HBV. We also describe the impact of nuclease treatment on sequencing efficiency.

## MATERIALS AND METHODS

### Probe design

We downloaded 4499 HBV non-recombinant whole genomes from the Hepatitis B Virus Database (hbvdb.lyon.inserm.fr/HBVdb/) that included the following number of samples of each genotype: A: 506, B: 1218, C: 1447, D: 823, E: 254, F: 197, G: 28 and H: 26. We used RaxML ^18^ with a general time reversible model with Gamma model of rate heterogeneity (“-m GTRGAMMA” option in RAxML) to infer a maximum likelihood phylogeny. We midpoint rooted the tree. Next, we used RAxML to infer the ancestral sequences and as input we used the midpoint rooted tree and the sequence of our isolates with the GTRGAMMA option. We used the ancestral sequence at the root of the tree to design the first set of probes assuming that this sequence on average has the least amount of divergence relative to all other isolates.

As the HBV genome is circular, we added 120 bases from the beginning of the ancestral root sequence to the end of the sequence to ensure that capture probes cover the break point which is used to present the genome linearly. We then divided the ancestral root sequence into 120 nt segments with 60 bases overlap which resulted in 55 probes. Genotype G has an insertion of 36 bases relative to other genotypes in the core gene. As the ancestral root sequence contained this insertion, we designed a probe of 120 nt which lacked this insertion. Furthermore, genotype D has a deletion of 33 bases in the pre-S1 region relative to all other genotypes. To ensure a probe covers this region for genotype D, we designed a probe of 120 nt that lacked this region.

Our previous work in HCV probe-based sequence capture demonstrated that probes of 120 bases long can tolerate up to 20% divergence relative to their target sequence before the efficiency of capture drops ^17^. To ensure that the designed probes are within 20% divergence of each of the isolates, we divided the isolates based on their genotype and created a consensus sequence for each genotype. We then aligned the probes to each isolate sequence and measured the proportion of mismatches between the probe and the isolate. For each isolate if a continuous region of at least 60 bases had probes which diverged from it more than 20%, a new probe was designed for the region using the genotype consensus sequence. As a quality control step, we removed any potential probe sequences that contained an “N” as we assumed that the sequence may be of low quality. Additionally we counted the number of ambiguous nucleotides and any sequence containing five or more ambiguous nucleotides was also removed. The final probe set contained 74 probes.

The probe sequences and the set of sequences that were used for their design are attached as supplementary material and can also be downloaded from the following webpage: https://doi.org/10.6084/m9.?gshare.22127015.

### Sample collection and preparation

#### (i) Clinical samples

We used plasma samples collected from adults with chronic HBV infection attending outpatient clinics at Oxford University Hospitals (OUH) NHS Foundation Trust. Approval for this work was provided by Oxford Research Ethics Committee A (ref. 09/H0604/20). Written informed consent for participation was provided by patients at enrollment.

Twenty plasma samples with viral loads <10,000 IU/mL were selected for this study. Viral loads were performed on the Abbott M2000 platform by OUH diagnostic microbiology laboratory (Table 1). Blood samples were collected in EDTA. To separate plasma, we centrifuged whole blood at 1800 rpm for 10 minutes. We removed the supernatant and stored it at -80 °C. Total nucleic acid was isolated from 1 mL plasma using the NucliSENS Easymag magnetic extraction system (bioMerieux) and collected in 50 μl of kit elution buffer for storage at –20 °C in aliquots (Figure 1B).

**Table 1:** Viral load and sequencing output for 20 plasma samples from adults with chronic HBV infection sequenced using an Illumina protocol for full-length HBV genome retrieval (± probe-based enrichment).

| Sample No. | Viral load (log10 IU/ml) | Subtype | Nuclease treatment | No capture |  |  |  |  |  | Capture |  |  |  |  |  |
| --- | --- | --- | --- | --- | --- | --- | --- | --- | --- | --- | --- | --- | --- | --- | --- |
|  |  |  |  | Read count | Reads pass-QC | HBV reads | HBV reads per million | % HBV genome assembled | Median coverage | Read count | Reads pass-QC | HBV reads | HBV reads per million | % HBV genome assembled | Median coverage |
| 1 | 4.07 | A2 | none | 20,933,044 | 17,463,756 | 0 | 0.0 | 0 | 0 | 15,053,030 | 12,226,942 | 6,322 | 420.0 | 95 | 9 |
|  |  |  | rnase | 14,270,072 | 10,029,188 | 2 | 0.1 | 0 | 0 | 9,023,230 | 5,671,256 | 17,702 | 1961.8 | 94 | 10 |
|  |  |  | dnase | 10,581,458 | 8,073,716 | 0 | 0.0 | 0 | 0 | 7,798,344 | 5,792,644 | 1,267 | 162.5 | 47 | 0 |
| 2 | 3.89 | A1 | none | 24,829,796 | 20,747,830 | 2 | 0.1 | 0 | 0 | 17,997,266 | 14,653,472 | 11,111 | 617.4 | 96 | 10 |
|  |  |  | rnase | 10,747,028 | 6,694,248 | 8 | 0.7 | 0 | 0 | 7,821,836 | 4,517,138 | 59,270 | 7577.5 | 98 | 21 |
|  |  |  | dnase | 15,739,300 | 12,156,038 | 0 | 0.0 | 0 | 0 | 11,355,572 | 8,337,602 | 559 | 49.2 | 38 | 0 |
| 3 | 3.52 | A | none | 21,145,796 | 17,416,474 | 0 | 0.0 | 0 | 0 | 15,114,326 | 12,080,868 | 393 | 26.0 | 48 | 0 |
|  |  |  | rnase | 9,412,374 | 5,663,302 | 0 | 0.0 | 0 | 0 | 6,343,938 | 3,216,884 | 361 | 56.9 | 33 | 0 |
|  |  |  | dnase | 10,173,230 | 7,778,512 | 0 | 0.0 | 0 | 0 | 7,706,602 | 5,777,898 | 0 | 0.0 | 0 | 0 |
| 4 | 3.36 | A | none | 20,230,320 | 17,013,454 | 0 | 0.0 | 0 | 0 | 14,408,928 | 11,803,456 | 2,085 | 144.7 | 88 | 3 |
|  |  |  | rnase | 10,919,638 | 6,437,256 | 0 | 0.0 | 0 | 0 | 7,411,744 | 3,744,028 | 4,837 | 652.6 | 79 | 3 |
|  |  |  | dnase | 18,696,910 | 15,029,678 | 0 | 0.0 | 0 | 0 | 13,755,642 | 10,784,290 | 655 | 47.6 | 53 | 1 |
| 5 | 3.19 | D1 | none | 25,862,232 | 21,354,586 | 0 | 0.0 | 0 | 0 | 19,474,444 | 15,652,348 | 401 | 20.6 | 48 | 0 |
|  |  |  | rnase | 9,415,412 | 4,746,086 | 0 | 0.0 | 0 | 0 | 6,972,594 | 2,999,110 | 2,274 | 326.1 | 64 | 1 |
|  |  |  | dnase | 21,208,452 | 17,124,384 | 0 | 0.0 | 0 | 0 | 15,019,534 | 11,643,926 | 19 | 1.3 | 10 | 0 |
| 6 | 3.18 | D5 | none | 19,398,596 | 16,257,044 | 0 | 0.0 | 0 | 0 | 13,052,120 | 10,503,632 | 88 | 6.7 | 0 | 0 |
|  |  |  | rnase | 11,050,832 | 6,153,260 | 0 | 0.0 | 0 | 0 | 8,240,228 | 4,077,122 | 1,430 | 173.5 | 58 | 1 |
|  |  |  | dnase | 11,156,224 | 9,533,464 | 0 | 0.0 | 0 | 0 | 7,665,006 | 6,311,486 | 0 | 0.0 | 0 | 0 |
| 7 | 2.89 | D1 | none | 21,290,168 | 18,053,908 | 0 | 0.0 | 0 | 0 | 15,201,562 | 12,546,846 | 410 | 27.0 | 26 | 0 |
|  |  |  | rnase | 6,604,634 | 3,277,524 | 0 | 0.0 | 0 | 0 | 4,881,446 | 2,156,000 | 2,430 | 497.8 | 64 | 1 |
|  |  |  | dnase | 12,435,072 | 10,610,356 | 0 | 0.0 | 0 | 0 | 8,532,044 | 7,076,998 | 0 | 0.0 | 0 | 0 |
| 8 | 2.88 | C2 | none | 17,752,464 | 14,917,026 | 0 | 0.0 | 0 | 0 | 12,927,836 | 10,557,494 | 1,171 | 90.6 | 59 | 1 |
|  |  |  | rnase | 11,440,204 | 7,762,324 | 0 | 0.0 | 0 | 0 | 7,256,430 | 4,453,350 | 12,093 | 1666.5 | 87 | 2 |
|  |  |  | dnase | 15,164,780 | 12,329,492 | 0 | 0.0 | 0 | 0 | 10,873,428 | 8,574,036 | 61 | 5.6 | 2 | 0 |
| 9 | 2.78 | B3 | none | 12,586,968 | 9,477,256 | 0 | 0.0 | 0 | 0 | 9,346,712 | 6,768,854 | 1,163 | 124.4 | 55 | 1 |
|  |  |  | rnase | 3,725,686 | 963,994 | 0 | 0.0 | 0 | 0 | 2,924,800 | 720,506 | 5,429 | 1856.2 | 61 | 1 |
|  |  |  | dnase | 3,391,284 | 1,846,784 | 0 | 0.0 | 0 | 0 | 2,578,902 | 1,319,130 | 0 | 0.0 | 0 | 0 |
| 10 | 2.77 | B2 | none | 23,567,564 | 17,763,880 | 0 | 0.0 | 0 | 0 | 17,390,788 | 12,429,772 | 168 | 9.7 | 34 | 0 |
|  |  |  | rnase | 10,374,034 | 5,927,086 | 0 | 0.0 | 0 | 0 | 7,406,372 | 3,658,894 | 893 | 120.6 | 26 | 0 |
|  |  |  | dnase | 14,029,578 | 10,515,512 | 0 | 0.0 | 0 | 0 | 10,342,508 | 7,400,774 | 0 | 0.0 | 0 | 0 |
| 11 | 2.76 | D3 | none | 14,323,558 | 12,356,090 | 0 | 0.0 | 0 | 0 | 9,752,656 | 8,196,682 | 1,041 | 106.7 | 0 | 0 |
|  |  |  | rnase | 7,208,416 | 3,938,548 | 0 | 0.0 | 0 | 0 | 5,259,304 | 2,551,594 | 3,922 | 745.7 | 54 | 1 |
|  |  |  | dnase | 5,729,868 | 4,617,560 | 0 | 0.0 | 0 | 0 | 3,941,876 | 3,038,332 | 134 | 34.0 | 0 | 0 |
| 12 | 2.76 | D1 | none | 18,172,348 | 15,403,362 | 0 | 0.0 | 0 | 0 | 12,879,278 | 10,591,494 | 308 | 23.9 | 0 | 0 |
|  |  |  | rnase | 11,409,822 | 7,606,280 | 0 | 0.0 | 0 | 0 | 7,586,406 | 4,563,632 | 1,497 | 197.3 | 20 | 0 |
|  |  |  | dnase | 9,211,228 | 7,223,996 | 0 | 0.0 | 0 | 0 | 6,681,556 | 5,032,218 | 0 | 0.0 | 0 | 0 |
| 13 | 2.68 | E | none | 23,259,426 | 19,678,708 | 0 | 0.0 | 0 | 0 | 16,773,174 | 13,860,314 | 805 | 48.0 | 73 | 1 |
|  |  |  | rnase | 5,733,202 | 1,918,436 | 0 | 0.0 | 0 | 0 | 4,619,986 | 1,327,688 | 3,566 | 771.9 | 78 | 2 |
|  |  |  | dnase | 24,115,084 | 21,506,066 | 0 | 0.0 | 0 | 0 | 17,199,360 | 15,054,188 | 88 | 5.1 | 0 | 0 |
| 14 | 2.64 | C2 | none | 19,579,800 | 15,530,932 | 0 | 0.0 | 0 | 0 | 13,960,764 | 10,578,950 | 1,261 | 90.3 | 54 | 1 |
|  |  |  | rnase | 6,203,356 | 2,795,282 | 0 | 0.0 | 0 | 0 | 4,790,500 | 1,942,126 | 4,706 | 982.4 | 70 | 2 |
|  |  |  | dnase | 13,742,552 | 11,692,072 | 0 | 0.0 | 0 | 0 | 9,554,018 | 7,908,836 | 358 | 37.5 | 0 | 0 |
| 15 | 2.55 | D3 | none | 18,320,118 | 14,323,370 | 0 | 0.0 | 0 | 0 | 13,272,736 | 9,921,674 | 30 | 2.3 | 0 | 0 |
|  |  |  | rnase | 11,177,644 | 6,480,840 | 0 | 0.0 | 0 | 0 | 7,758,056 | 3,926,498 | 316 | 40.7 | 6 | 0 |
|  |  |  | dnase | 9,910,562 | 7,166,388 | 0 | 0.0 | 0 | 0 | 7,208,138 | 4,939,854 | 0 | 0.0 | 0 | 0 |
| 16 | 2.52 | D2 | none | 17,431,576 | 14,992,498 | 0 | 0.0 | 0 | 0 | 12,095,246 | 10,120,378 | 1,037 | 85.7 | 35 | 0 |
|  |  |  | rnase | 7,588,514 | 3,608,292 | 0 | 0.0 | 0 | 0 | 5,724,212 | 2,401,742 | 3,292 | 575.1 | 58 | 1 |
|  |  |  | dnase | 9,681,050 | 7,598,656 | 0 | 0.0 | 0 | 0 | 6,860,588 | 5,180,184 | 92 | 13.4 | 0 | 0 |
| 17 | 2.49 | D2 | none | 13,876,540 | 10,698,164 | 0 | 0.0 | 0 | 0 | 9,820,456 | 7,184,016 | 712 | 72.5 | 0 | 0 |
|  |  |  | rnase | 4,614,834 | 2,214,120 | 0 | 0.0 | 0 | 0 | 3,440,026 | 1,469,952 | 1,275 | 370.6 | 43 | 0 |
|  |  |  | dnase | 3,826,436 | 2,028,754 | 0 | 0.0 | 0 | 0 | 2,420,374 | 1,219,428 | 0 | 0.0 | 0 | 0 |
| 18 | 2.45 | E | none | 20,962,554 | 17,802,228 | 0 | 0.0 | 0 | 0 | 15,212,438 | 12,532,142 | 0 | 0.0 | 0 | 0 |
|  |  |  | rnase | 10,444,192 | 5,771,954 | 0 | 0.0 | 0 | 0 | 7,767,042 | 3,705,066 | 291 | 37.5 | 23 | 0 |
|  |  |  | dnase | 20,362,950 | 17,646,582 | 0 | 0.0 | 0 | 0 | 14,668,534 | 12,452,388 | 0 | 0.0 | 0 | 0 |
| 19 | 2.02 | A | none | 22,997,532 | 18,572,882 | 0 | 0.0 | 0 | 0 | 17,012,832 | 13,351,474 | 120 | 7.1 | 0 | 0 |
|  |  |  | rnase | 14,307,796 | 9,990,204 | 0 | 0.0 | 0 | 0 | 9,369,470 | 5,866,900 | 157 | 16.8 | 0 | 0 |
|  |  |  | dnase | 8,312,318 | 6,056,304 | 0 | 0.0 | 0 | 0 | 6,610,638 | 4,703,870 | 0 | 0.0 | 0 | 0 |
| 20 | 1.98 | A5 | none | 18,542,206 | 15,610,616 | 0 | 0.0 | 0 | 0 | 13,399,040 | 11,001,066 | 63 | 4.7 | 12 | 0 |
|  |  |  | rnase | 6,103,116 | 3,121,634 | 0 | 0.0 | 0 | 0 | 4,642,700 | 2,144,522 | 928 | 199.9 | 23 | 0 |
|  |  |  | dnase | 9,481,756 | 6,000,322 | 0 | 0.0 | 0 | 0 | 7,597,368 | 4,648,590 | 0 | 0.0 | 0 | 0 |

**Figure 1:**
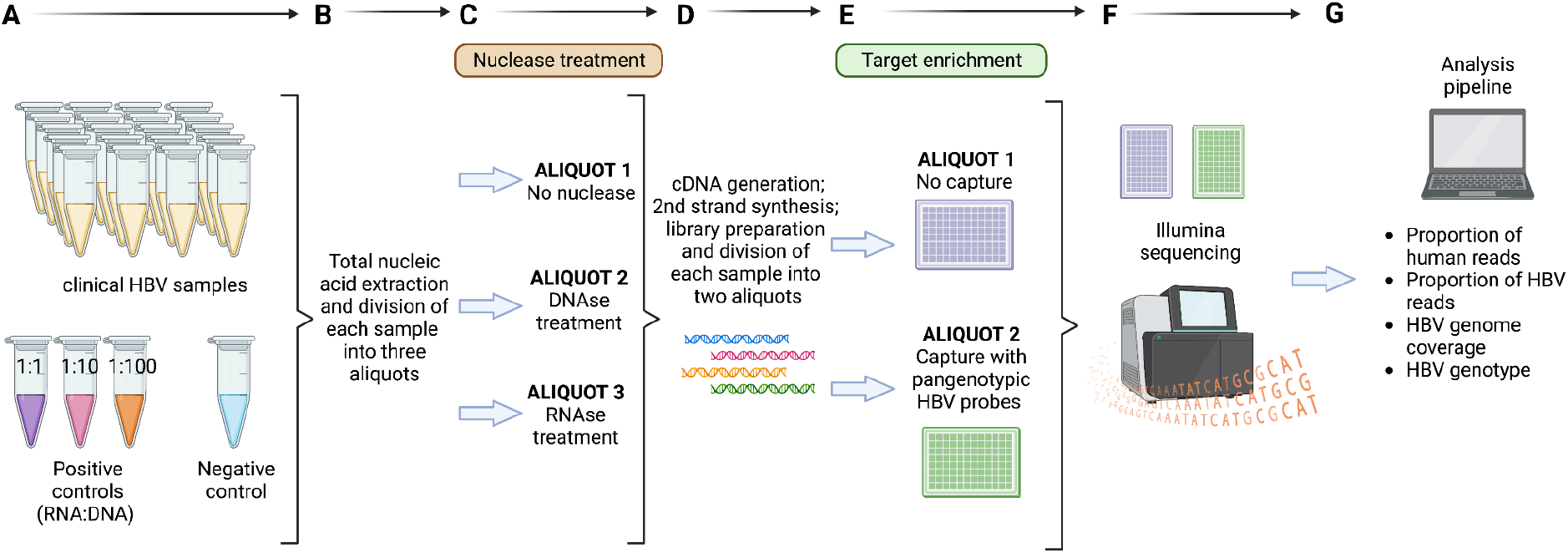
Schematic diagram showing HBV sequencing workflow. A: Sample and control description; B: Nucleic acid extraction; C: Nuclease treatment; D: Library prep; E: Probe-based enrichment (capture); F: Illumina sequencing; G: Bioinformatic analysis.

#### (ii) Control samples

Human Total Brain RNA (ThermoFisher Scientific) and a 48,502 bp double-stranded linear DNA from Bacteriophage Lambda (ThermoFisher Scientific) were used. Both were quantified with Qubit dsDNA High Sensitivity and RNA High Sensitivity Assay kits and mixed to create control samples of known RNA:DNA ratios (1:1, 1:10, 1:100).

All samples, including clinical and controls, were quantified using Qubit dsDNA High Sensitivity and RNA High Sensitivity Assay kits (ThermoFisher Scientific) and stored in aliquots at -20°C.

### Nuclease treatment

Each of the 24 samples (20 clinical and 4 control samples) were processed in triplicate as follows:

#### (i) No nuclease treatment

Nuclease free water was used to make the sample up to a final volume of 50μL.

#### (ii) DNAse treatment

DNA was depleted using TURBO DNase (ThermoFisher Scientific) as per manufacturer’s instructions. Briefly, a reaction containing 12 μL of sample material, 1 µL of TURBO DNase (2 U), 5 µL of 10x TURBO DNase Buffer and 32 µL of nuclease free water was incubated at 37 °C for 30 min. The reaction was inactivated using 1.5 μL EDTA (0.5M) (ThermoFisher) followed by incubation at 75 °C for 10 min.

#### (iii) RNAse treatment

12 μL of sample/control was denatured at 94 °C for 2 min and snap-frozen on ice. RNA digestion was performed with 0.5 µL RNase A (NEB), 2.5 µL 1M TrisHcl (ThermoFisher) and 35 µL of nuclease free water, followed by incubation at 56 °C for 5 min followed by snap-chilling. The reaction was inactivated at 25 °C for 10 min with 1.25 μL of Murine RNAse inhibitor (New England Biolabs).

### Library Preparation

Libraries were prepared for Illumina sequencing using the Takara Bio SMARTer Stranded Total RNA-Seq Kit v3 - Pico Input Mammalian (Fig 1D) with the following minor adjustments to the manufacturer’s recommendations. Sample volumes were concentrated to 3µl using SPRIselect beads (Beckman Coulter) and processed using a 0.25x reduced scale miniaturised reaction volume for first strand cDNA synthesis. Indexing PCR was completed using Oxford Genomic Consortium’s in-house indexed primers and 12 amplification cycles. Fragment size distribution of the final libraries were analysed on the Agilent 4200 TapeStation using High Sensitivity D1000 DNA screentape. The amplified cDNA libraries corresponding to each aliquot were pooled in equivolume proportions to generate a final multiplex library. The pool was purified using SPRIselect beads and subsequently quantified using High Sensitivity dsDNA Qubit assay (Invitrogen) and re-analysed using D1000 DNA screentape (Agilent).

### Hybrid capture of sequencing libraries

A 4.4µg aliquot of the final multiplexed library was enriched for HBV using the custom-designed probe panel and xGen Hybridization and Wash Kit (IDT Technologies) following manufacturer’s instructions. The final enriched library was amplified (12 cycles on-bead PCR), repurified and normalised, then sequenced on the NovaSeq6000 platform with 2×150nt paired-end reads.

### Metagenomic sequencing

An aliquot of the metagenomic pool was normalised to 10nM and sequenced on a NovaSeq6000 instrument with a 2×150nt paired-end read length.

### Bioinformatic processing

De-multiplexed sequence read-pairs were trimmed of low-quality bases using QUASR ^19^ and adapter sequences with CutAdapt ^20^ and Skewer ^21^ and subsequently discarded if either read had less than 50bp sequence remaining. The cleaned read pairs were mapped to human reference genome hg19 using Bowtie ^22^ and excluded from further analyses. All nonhuman read pairs were mapped using BWA-MEM^23^ to a set of 44 HBV references covering all known HBV genotypes and subgenotypes to choose an appropriate reference ^24^. The HBV reference with the most number of HBV reads mapping to it was chosen as the genetically closest reference to the sequenced isolate. Next, all nonhuman read pairs were mapped to the closest HBV reference. Picard markduplicates tool (http://broadinstitute.github.io/picard) was then used to remove duplicate read pairs (where read pairs starting in the same place and ending in the same place on the genome are assumed to be PCR duplicates).

## RESULTS

### Probe-based enrichment increases HBV sequence yield

We compared the sequencing output of pooled libraries of 20 samples with/without HBV specific probe-based enrichment using a panel of biotinylated DNA oligonucleotides (termed “capture” vs “no-capture” respectively), samples were sequenced in triplicate (under varying nuclease pre-treatments, discussed later). Sample HBV viral loads ranged from 1.98 log_10_ to 4.07 log_10_, representing harder to sequence low viral loads, and included samples from genotypes A, B, C, D and E (Table 1).

Capture increased the number of HBV reads per million reads sequenced in all genotypes. In the “no nuclease” arem, capture resulted in a 47,970 fold increase in the number of reads mapped to the HBV genome (243 HBV reads per million reads sequenced in the capture pool vs. 5×10^−3^ reads per million in the no-capture pool). Out of 20 samples, only 1/20 without capture generated HBV reads (viral load 3.89 log_10_ IU/ml), this increased to 19/20 samples with capture. HBV sequence yield was increased in the capture arm and resulted in 2.30 log_10_ (95% confidence interval 1.99 - 2.48 log_10_) increase in HBV reads (per million reads sequenced) per log_10_ increase in viral load (Figure 2) when a linear relationship was assumed. Capture increased the proportion of HBV reads from a median of 0 (range 0 - 8.05 ×10^−8^) to a median of 3.75 ×10^−5^ (range 0 - 6.17×10^−4^) whilst decreasing the proportion of human reads (Supplementary figure 1).

**Figure 2:**
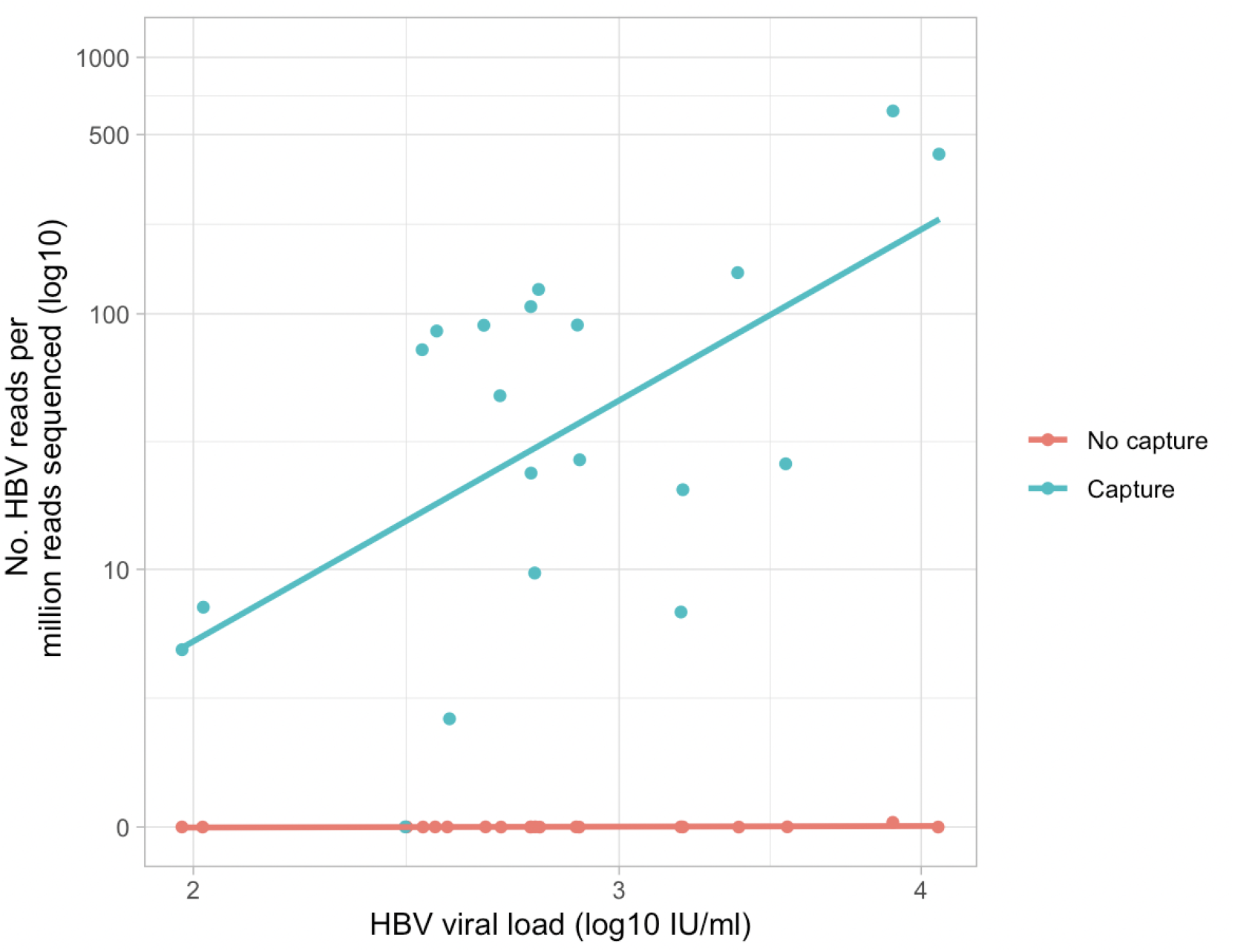
HBV specific probe-based capture increased the efficiency of HBV sequencing by nearly 48,000 fold. Assuming a linear relationship between sequence yield and viral load, we observed a 2.30 log_10_ increase in the number of HBV reads per log_10_ increase in HBV DNA viral load in plasma.

### RNase and DNase treatment impact on sequencing output

We compared the impact of RNAse and DNAse treatment after extraction on sequencing output on samples undergoing probe-based enrichment. First, the enzyme treatments were trialed on control material to assess the depletion efficacy. Three control samples were prepared containing different DNA: RNA ratios; 1:1, 10:1 and 100:1, using Total brain RNA and Lambda bacteriophage DNA. RNAse treatment effectively depleted the RNA control reducing the number of reads mapping to the human genome. DNAse treatment effectively depleted the DNA control reducing reads mapping to the DNA control genome (Lambda bacteriophage) (Supplementary figure 2).

We then explored the effect of RNAse and DNAse treatments of HBV infected clinical samples undergoing probe-based enrichment. RNAse treatment increased the number of HBV reads per million reads sequenced, whereas DNAse treatment decreased the number of HBV reads (per million reads sequenced, Figure 3). The proportion of HBV reads increased a median of 12 fold with RNAse treatment from a median of 3.75×10^−5^ (range 0 - 6×10^−4^) to 4.34×10^−4^ (range 1.68×10^−5^ - 7.6×10^−3^), but decreased with DNAse treatment to a median of 0 (range 0 - 1.6×10^−4^). A similar trend where RNAse treatment improves DNA sequencing was observed for the controls (Supplementary figure 3). Among the HBV infected sampled the proportion of human reads decreased with RNAse treatment, from a median of 0.52 (range 0.42 - 0.67) to 0.34 (range 0.11 - 0.57), but remained static with DNAse treatment with a median of 0.51 (range 0.28 - 0.69) (Supplementary figure 4).

**Figure 3:**
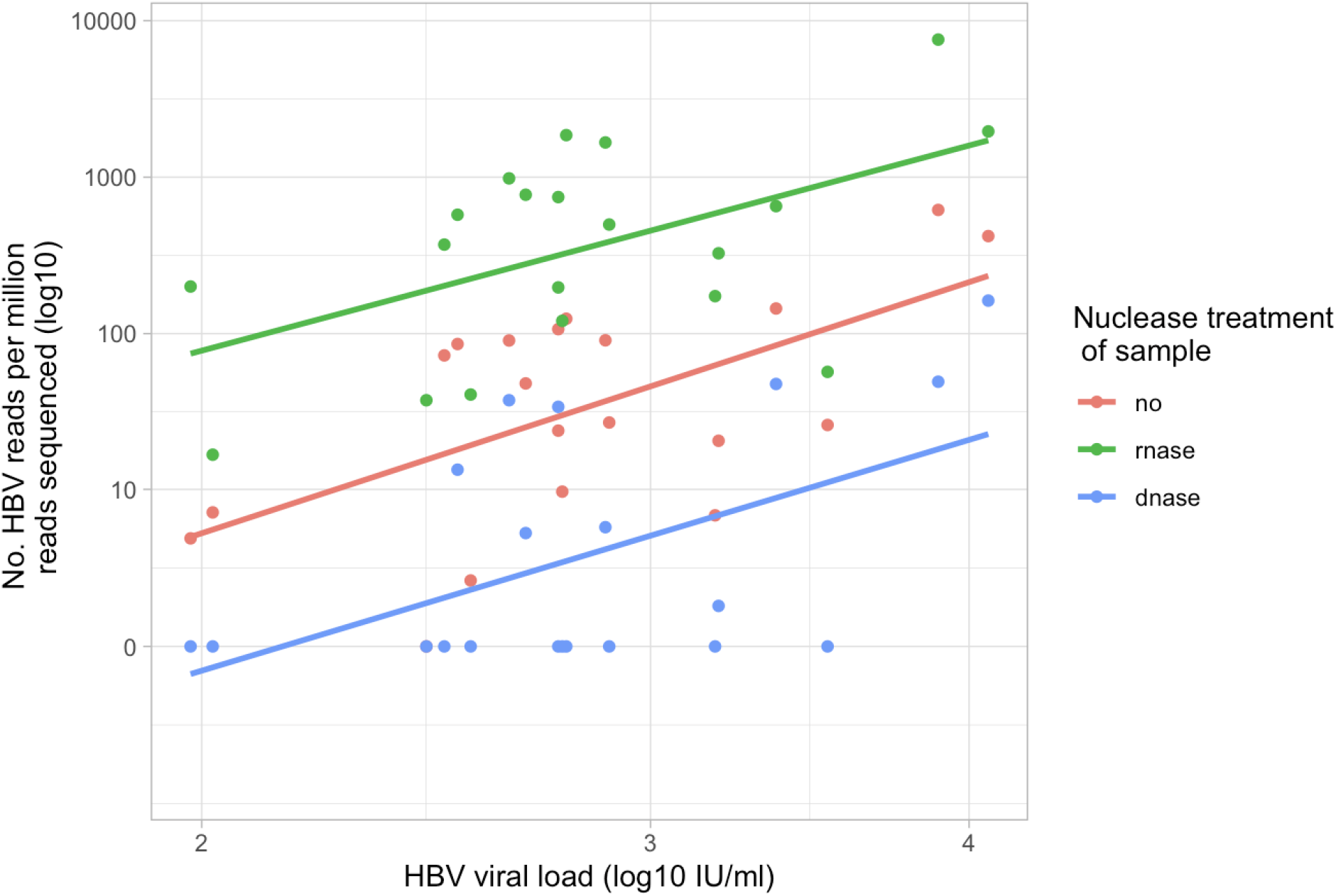
Impact of RNAse and DNAse treatment prior to library prep on the proportion of reads mapping to the HBV genome in 20 clinical samples.

Median depth of coverage of the HBV genome was low in this study (Table 1) due to the low viral loads. A comparison of percentage genome assembled for different nuclease treatments is shown in Supplementary figure 5a. Depth of coverage was variable across the genome (see Supplementary figure 5b for an example coverage plot for a sample with viral load 3.89 log_10_ IU/ml). Peaks of higher coverage were seen around positions 1600-2000 with lower coverage seen between positions 650-1500.

## DISCUSSION

HBV sequencing is key to informing questions pertinent to epidemiology, transmission, persistence, disease outcomes and treatment/vaccine responsiveness. To date, this field is hampered by poor investment and inadequate approaches to deal with low viral load samples, which are common in clinical practice. We developed a targeted sequencing method using a custom panel of biotinylated oligos that increases the sequencing efficacy of HBV. Capture increased the number of samples with HBV reads from 1/20 to 19/20, increased the proportion of HBV reads and percentage genome coverage, whilst decreasing the proportion of human reads. The probe set is designed to be pan-genotypic; here sequencing of genotypes A, B, C, D and E is demonstrated, accounting for >95% of the global burden of HBV ^25^.

HBV is a dsDNA virus that replicates via an RNA intermediate. Since the Takara Bio kit uses cDNA synthesis and 2nd strand synthesis steps to convert RNA to DNA prior to performing a DNA library prep and Illumina sequencing, in theory both HBV rc-DNA and pgRNA could be contributing to sequencing output. The comparison of nuclease treatments allowed us to investigate this further. DNAse treatment reduced the number of HBV reads, completely eliminating HBV reads in 11/20 samples, whereas RNAse treatment improved sequencing, therefore it is likely that this method predominantly sequences HBV DNA rather than RNA.

The observation that HBV DNA rather than RNA is predominantly being sequenced seems paradoxical as the Takara Bio SMARTer Stranded Total RNA-Seq method was developed to sequence RNA targets. In theory, although RT preferentially targets an RNA template during cDNA synthesis, it is also capable of reading DNA templates. We hypothesise that when both DNA and RNA are present, RNA is preferentially processed, reducing the proportion of DNA converted to library. By removing the preferred template type (RNA) with RNAse treatment, more ssDNA is converted to dsDNA and therefore is available for library prep, and less reverse transcribed human RNA is present as background, improving the ratio of viral:host nucleic acid. Furthermore, the RNAse treatment protocol involves a denaturation step (94°C for 2 minutes followed by snap chilling), which may increase the availability of DNA for library prep.

This work forms the basis for more extensive characterization of the impact of capture probes on HBV sequencing output. The viral loads of samples here (1.98 log_10_ to 4.07 log_10_ IU/ml) clustered at the lower end of the clinical range ^4^, therefore further experimental work is required to determine the impact of capture on higher viral load samples. Capture allowed HBV reads to be sequenced in 19/20 samples (vs. 1/20 without capture), however depth of coverage was variable across the genome. We saw a peak in depth of coverage between positions 1600-2000 a region encoding protein X, containing the 5’ end of the positive strand, the nick/overlap of the 3’ and 5’ ends of the negative strand, including direct repeats 1 and 2 (DR1, DR2). Coverage depth was low between positions 650-1500, this is the single stranded region of the genome. Sequencing strategies involving completion/ligation during which uses DNA polymerase and DNA ligase turn the partially double stranded genome into a fully double stranded circle, could be used to tackle this issue.

Probe inefficiency at lower viral loads has also been observed elsewhere ^12^. Alternative methods with additional host depletion (e.g. benzonase or MNAse prior to extraction) or an amplification step prior to library prep e.g. amplicon based PCR) are likely to be required to reliably obtain full genomes in samples with viral loads <10^4^ IU/ml. In order to infer the consensus genome sequence of a sample accurately, a minimum of x5 coverage is required, but ideally the minimum coverage would be higher (x10-20).

Pan-genotype probe based enrichment can be used to improve the efficiency of viral sequencing by increasing the ratio of viral:human reads. However further method development is required to obtain whole genome sequences from low viral load samples.

## SUPPLEMENTARY FIGURES

**Supplementary figure 1.**
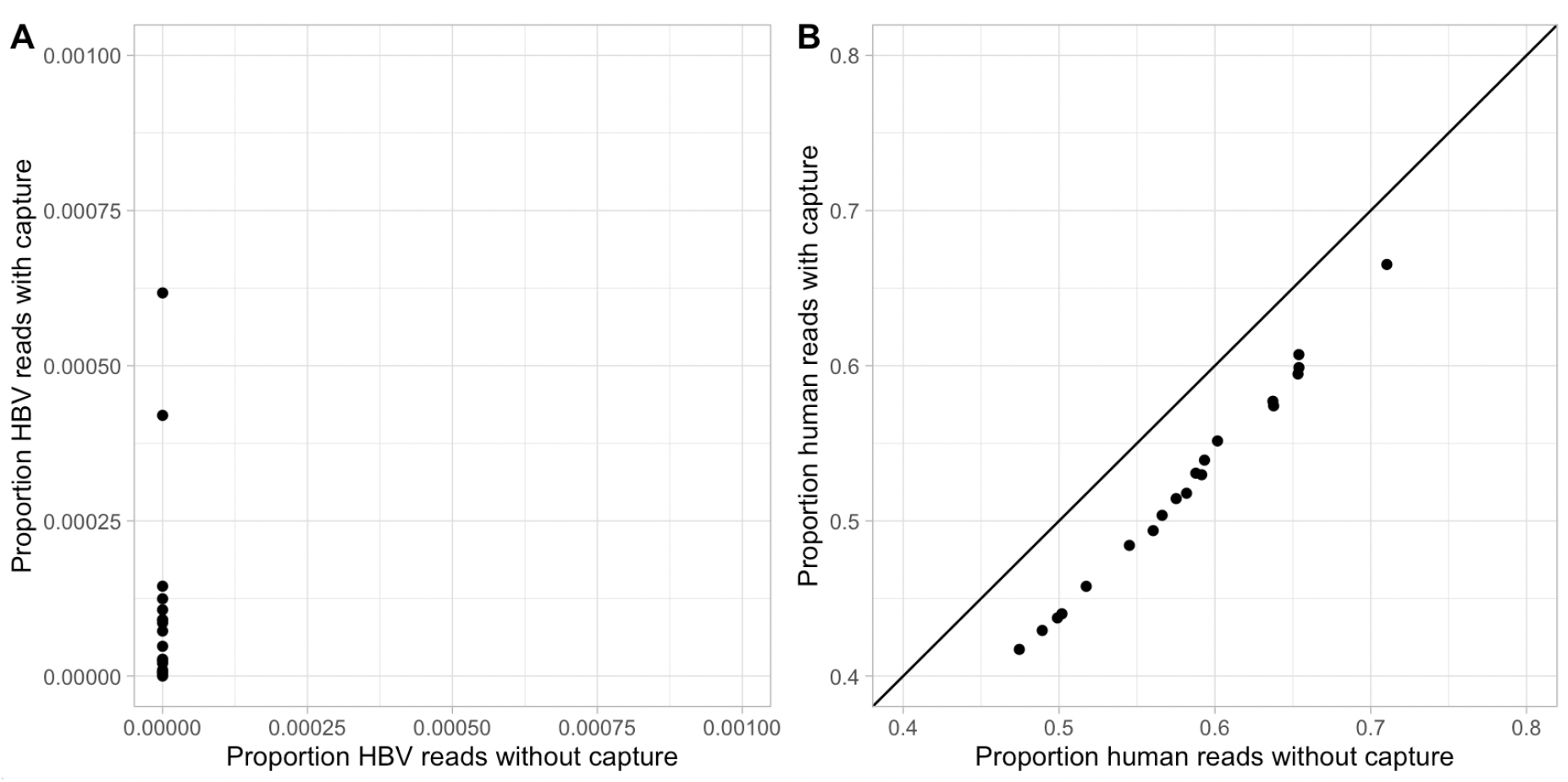
Impact of capture on proportion of reads mapping to a) HBV and b) human genome.

**Supplementary figure 2.**
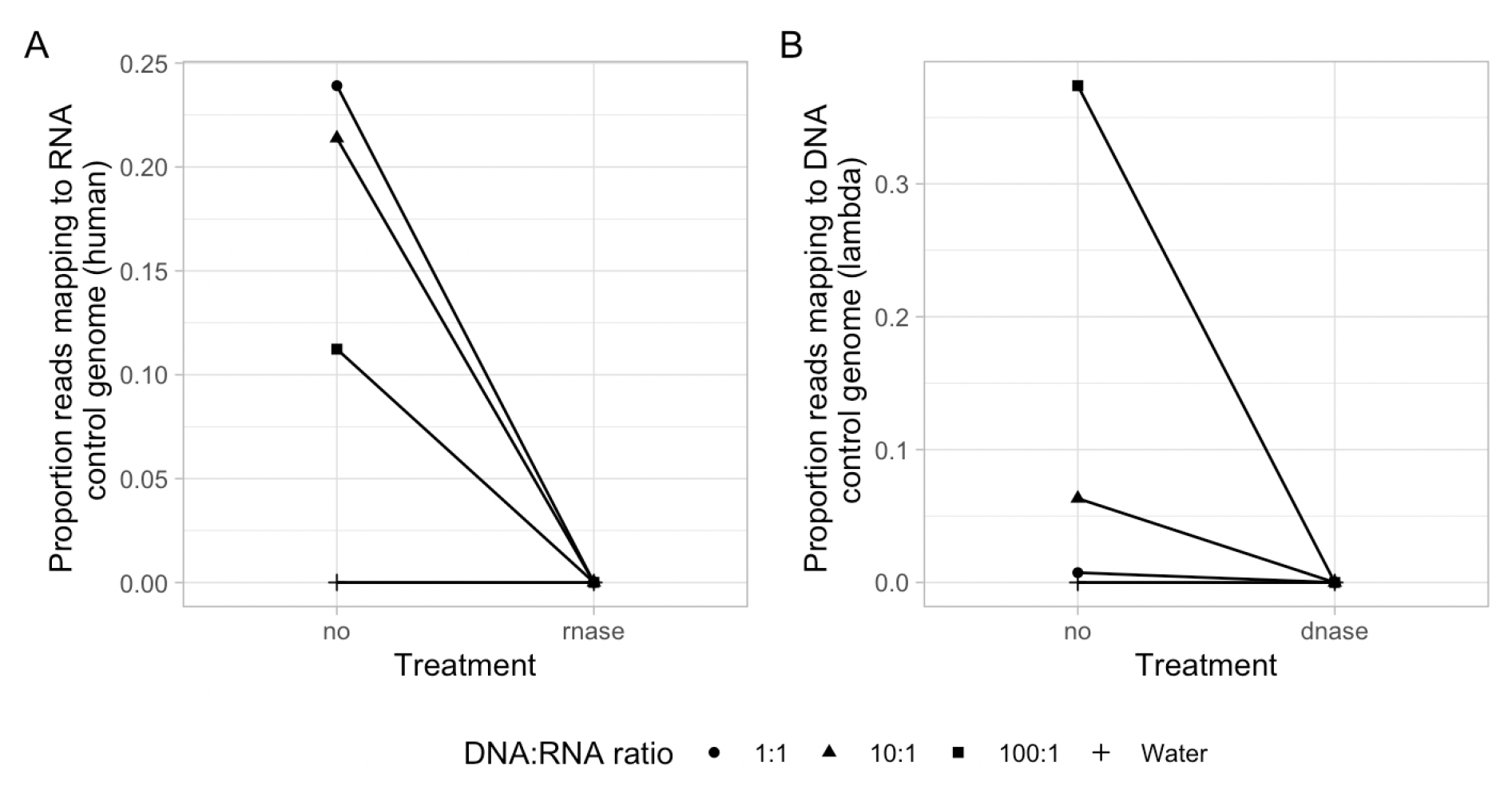
Depletion efficacy of nuclease treatments on control material. A. Proportion of reads mapping to RNA control genome following RNase treatment, B. Proportion of reads mapping to DNA control genome following DNase treatment

**Supplementary figure 3.**
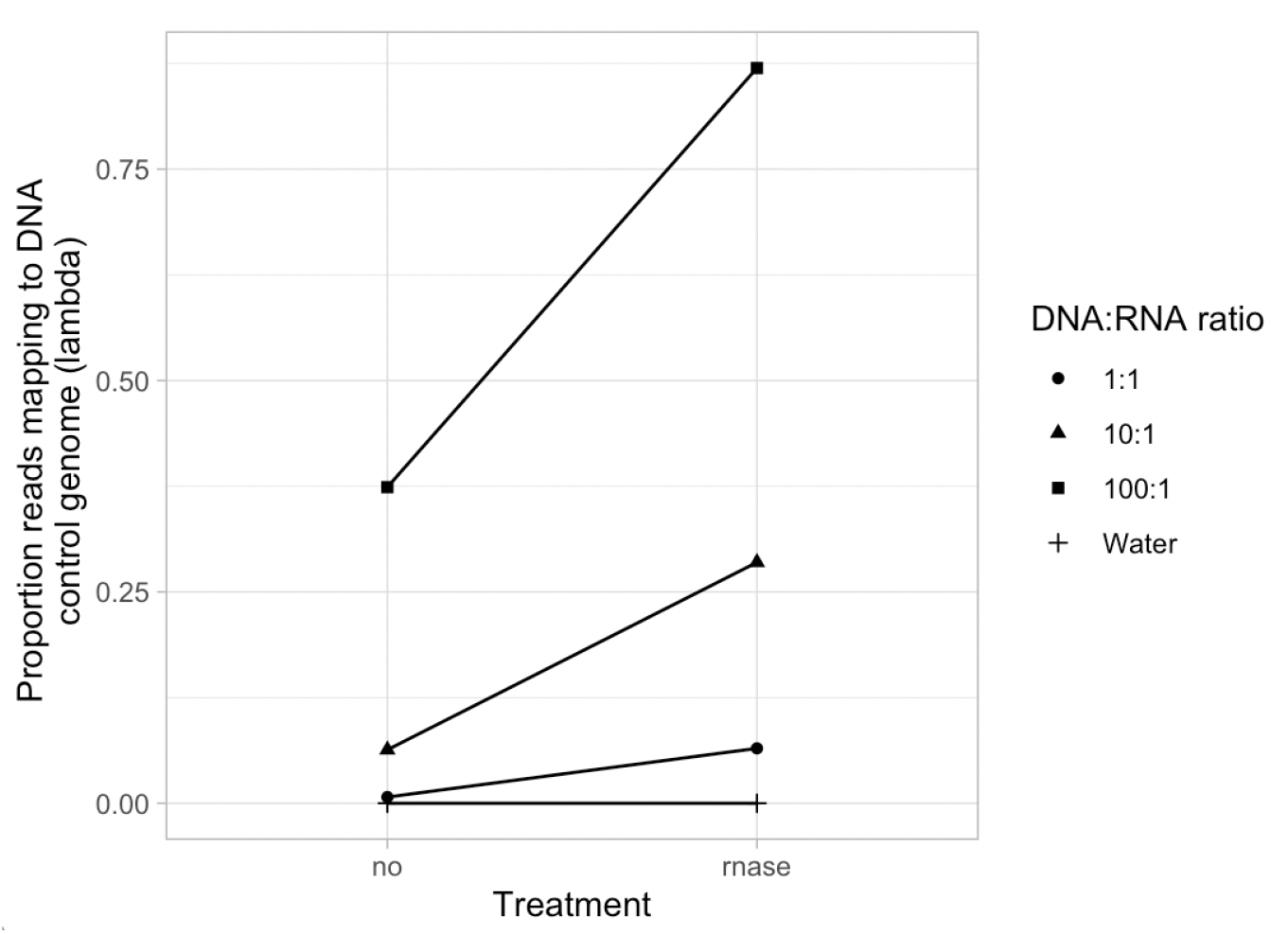
RNAse depletion protocol increases proportion of reads mapping to DNA control.

**Supplementary figure 4.**
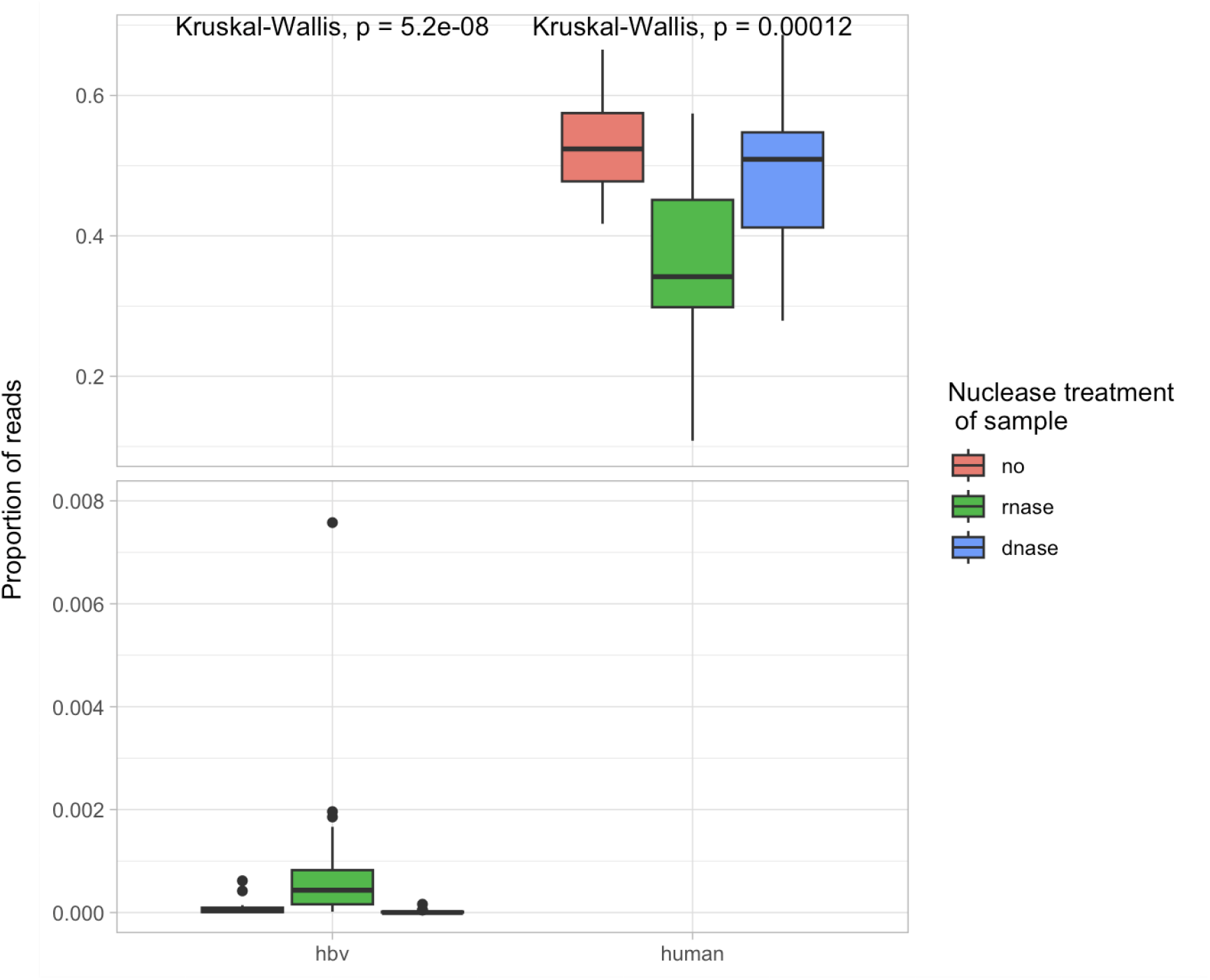
Impact of nuclease treatments on sequencing output of 20 HBV infected plasma samples. Proportion of reads mapping to HBV vs. human genomes.

**Supplementary figure 5.**
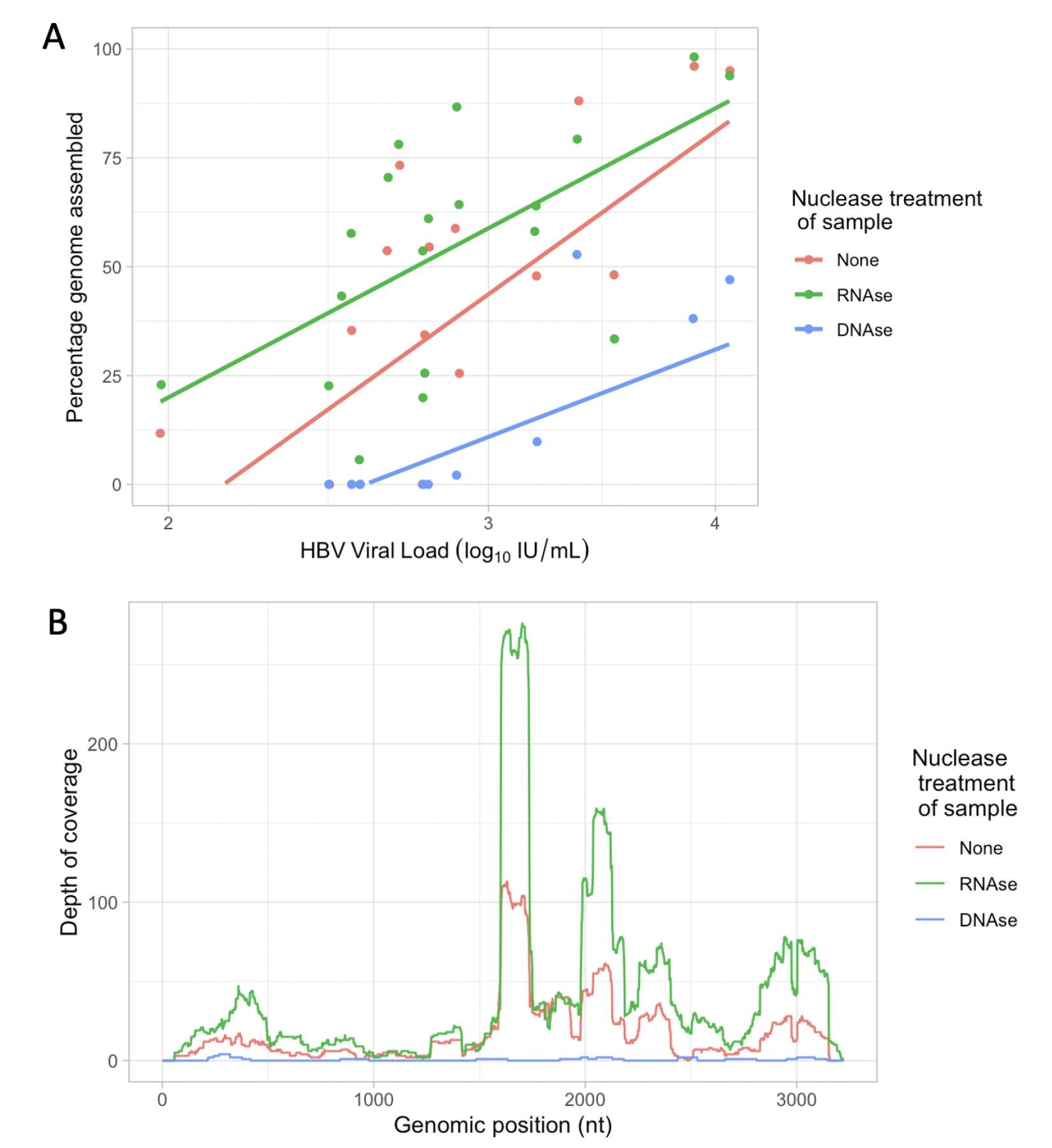
Impact of nuclease treatment on genome coverage. A. Impact of nuclease treatment on percentage genome assembled B. Example coverage plots for a sample with viral load 3.89 log_10_ under different nuclease treatment

## REFERENCES

1. GBD 2019 Hepatitis B Collaborators. Global, regional, and national burden of hepatitis B, 1990-2019: a systematic analysis for the Global Burden of Disease Study 2019. Lancet Gastroenterol Hepatol 7, 796–829 (2022).

2. Revill, P. A. et al. The evolution and clinical impact of hepatitis B virus genome diversity. Nat. Rev. Gastroenterol. Hepatol. 17, 618–634 (2020).

3. McNaughton, A. L. et al. Insights From Deep Sequencing of the HBV Genome-Unique, Tiny, and Misunderstood. Gastroenterology 156, 384–399 (2019).

4. Downs, L. O. et al. Bimodal distribution and set point HBV DNA viral loads in chronic infection: retrospective analysis of cohorts from the UK and South Africa. Wellcome Open Res 5, 113 (2020).

5. Conceição-Neto, N. et al. Modular approach to customise sample preparation procedures for viral metagenomics: a reproducible protocol for virome analysis. Sci. Rep. 5, 16532 (2015).

6. Günther, S. et al. A novel method for efficient amplification of whole hepatitis B virus genomes permits rapid functional analysis and reveals deletion mutants in immunosuppressed patients. J. Virol. 69, 5437–5444 (1995).

7. Wang, X. et al. Integrating nested PCR with high-throughput sequencing to characterize mutations of HBV genome in low viral load samples. Medicine 96, e7588 (2017).

8. McNaughton, A. L. et al. Illumina and Nanopore methods for whole genome sequencing of hepatitis B virus (HBV). Sci. Rep. 9, 7081 (2019).

9. Martel, N., Gomes, S. A., Chemin, I., Trépo, C. & Kay, A. Improved rolling circle amplification (RCA) of hepatitis B virus (HBV) relaxed-circular serum DNA (RC-DNA). J. Virol. Methods 193, 653–659 (2013).

10. Goldsmith, C. et al. Cas9-targeted nanopore sequencing reveals epigenetic heterogeneity after de novo assembly of native full-length hepatitis B virus genomes. Microbial Genomics vol. 7 Preprint at https://doi.org/10.1099/mgen.0.000507 (2021).

11. Gu, W. et al. Depletion of Abundant Sequences by Hybridization (DASH): using Cas9 to remove unwanted high-abundance species in sequencing libraries and molecular counting applications. Genome Biol. 17, 41 (2016).

12. Berg, M. G. et al. Advanced molecular surveillance approaches for characterization of blood borne hepatitis viruses. PLoS One 15, e0236046 (2020).

13. Yamaguchi, J. et al. Universal Target Capture of HIV Sequences From NGS Libraries. Frontiers in Microbiology vol. 9 Preprint at https://doi.org/10.3389/fmicb.2018.02150 (2018).

14. Briese, T. et al. Virome Capture Sequencing Enables Sensitive Viral Diagnosis and Comprehensive Virome Analysis. MBio 6, e01491–15 (2015).

15. Matranga, C. B. et al. Enhanced methods for unbiased deep sequencing of Lassa and Ebola RNA viruses from clinical and biological samples. Genome Biol. 15, 519 (2014).

16. Depledge, D. P. et al. Specific Capture and Whole-Genome Sequencing of Viruses from Clinical Samples. PLoS ONE vol. 6 e27805 Preprint at https://doi.org/10.1371/journal.pone.0027805 (2011).

17. Bonsall, D. et al. ve-SEQ: Robust, unbiased enrichment for streamlined detection and whole-genome sequencing of HCV and other highly diverse pathogens. F1000Res. 4, 1062 (2015).

18. Stamatakis, A. RAxML version 8: a tool for phylogenetic analysis and post-analysis of large phylogenies. Bioinformatics 30, 1312–1313 (2014).

19. Gaidatzis, D., Lerch, A., Hahne, F. & Stadler, M. B. QuasR: quantification and annotation of short reads in R. Bioinformatics 31, 1130–1132 (2015).

20. Martin, M. Cutadapt removes adapter sequences from high-throughput sequencing reads. EMBnet.journal 17, 10–12 (2011).

21. Jiang, H., Lei, R., Ding, S.-W. & Zhu, S. Skewer: a fast and accurate adapter trimmer for next-generation sequencing paired-end reads. BMC Bioinformatics 15, 182 (2014).

22. Langmead, B., Trapnell, C., Pop, M. & Salzberg, S. L. Ultrafast and memory-efficient alignment of short DNA sequences to the human genome. Genome Biol. 10, R25 (2009).

23. Li, H. & Durbin, R. Fast and accurate short read alignment with Burrows-Wheeler transform. Bioinformatics 25, 1754–1760 (2009).

24. McNaughton, A. L., Revill, P. A., Littlejohn, M., Matthews, P. C. & Ansari, M. A. Analysis of genomic-length HBV sequences to determine genotype and subgenotype reference sequences. J. Gen. Virol. 101, 271–283 (2020).

25. Velkov, S., Ott, J. J., Protzer, U. & Michler, T. The Global Hepatitis B Virus Genotype Distribution Approximated from Available Genotyping Data. Genes 9, (2018).

